# Diminished dynamic cerebral autoregulatory capacity with forced oscillations in mean arterial pressure with elevated cardiorespiratory fitness

**DOI:** 10.1101/190405

**Authors:** Lawrence Labrecque, Kevan Rahimaly, Sarah Imhoff, Myriam Paquette, Olivier Le Blanc, Simon Malenfant, Samuel JE Lucas, Damian M. Bailey, Jonathan D. Smirl, Patrice Brassard

**Affiliations:** Department of Kinesiology, Faculty of Medicine, Université Laval, Québec, Canada; Research center of the Institut universitaire de cardiologie et de pneumologie de Québec, Québec, Canada; School of Sport, Exercise and Rehabilitation Sciences, University of Birmingham, Birmingham, UK; Department of Physiology, University of Otago, Dunedin, New Zealand; Neurovascular Research. Laboratory, Faculty of Life Sciences and Education, University of South Wales, South Wales, UK; Faculty of Medicine, Reichwald Health Sciences Centre, University of British Columbia–Okanagan, Kelowna, BC, Canada; Health and Exercise Sciences, University of British Columbia Okanagan, Kelowna BC, Canada

**Keywords:** cardiorespiratory fitness, dynamic cerebral autoregulation, cerebral pressure-flow relationship

## Abstract

The effect that cardiorespiratory fitness has on the dynamic cerebral autoregulatory capacity during changes in mean arterial pressure (MAP) remains equivocal. Using a multiple-metrics approach, challenging MAP across the spectrum of physiological extremes (i.e. spontaneous through forced MAP oscillations), we characterized dynamic cerebral autoregulatory capacity in 19 male endurance athletes and 8 controls via three methods: [1] onset of regulation (i.e., time delay before an increase in middle cerebral artery (MCA) conductance [MCA flow velocity (MCAv)/mean arterial pressure (MAP)] and rate of regulation, after transient hypotension induced by sit-to-stand, and transfer function analysis (TFA) of MAP and MCAv responses during [2] spontaneous and [3] forced oscillations (5-min of squat-stand maneuvers performed at 0.05 and 0.10 Hz). Reductions in MAP and mean MCAv (MCAV_mean_) during initial orthostatic stress (0-30 s after sit-to-stand) and the prevalence of orthostatic hypotension were also determined. Onset of regulation was delayed after sit-to-stand in athletes (3.1±1.7 vs. 1.5±1.0 s; p=0.03), but rate of regulation was not different between groups (0.24±0.05 vs. 0.21±0.09 s^-1^; p=0.82). While both groups had comparable TFA metrics during spontaneous oscillations, athletes had higher TFA gain during 0.10 Hz squat-stand vs. recreational controls (p=0.01). Reductions in MAP (p=0.15) and MCAV_mean_ (p=0.11) during orthostatic stress and the prevalence of initial orthostatic hypotension (p=0.65) were comparable between groups. These results indicate an intact ability of the cerebral vasculature to react to spontaneous oscillations but an attenuated capability to counter rapid and large changes in MAP in individuals with elevated cardiorespiratory fitness.

**New findings:** *What is the central question of this study?:*

- What is the impact of cardiorespiratory fitness on dynamic cerebral autoregulatory capacity?

*What is the main finding and its importance?:*

- Our results indicate an intact ability of the cerebral vasculature to react to spontaneous oscillations but an attenuated capability to counter rapid and large changes in mean arterial pressure in individuals with elevated cardiorespiratory fitness.

## INTRODUCTION

Regular aerobic-endurance exercise has been associated with higher resting cerebral blood velocity in men (Ainslie *et al*., 2008), while higher cardiorespiratory fitness (CRF) has been shown to improve cerebral blood velocity during exercise (Brugniaux *et al*., 2014), as well as cerebrovascular reactivity to carbon dioxide (Bailey *et al*., 2013b; Barnes *et al*., 2013). However, whether CRF positively influences the ability of the cerebrovasculature to respond to rapid changes in mean arterial pressure (MAP) (traditionally referred to as dynamic cerebral autoregulation) remains equivocal. A weakened dynamic cerebral autoregulatory capacity, characterized by a delayed onset of regulation following transient hypotension induced by thigh-cuff deflation and larger normalized transfer function gain of spontaneous oscillations in MAP and flow velocity in the middle cerebral artery (MCAv), has been reported in endurance athletes compared to sedentary controls (Lind-Holst *et al*., 2011). Conversely, comparable dynamic cerebral autoregulatory capacity between endurance athletes and sedentary controls has also been reported following thigh-cuff deflation [rate of regulation (RoR)] and via transfer function analysis (TFA; coherence – fraction of the MAP which is linearly related to MCAv, gain – amplitude of MCAv change for a given oscillation in MAP, phase – difference in the timing of the MAP and MCAv waveforms) of spontaneous or forced oscillations in MAP and MCAv in young and elderly athletes (Aengevaeren *et al*., 2013; Ichikawa *et al*., 2013). Taking into consideration the notion that most metrics are unrelated to each other (Tzeng *et al*., 2012), direct comparison between studies using different analysis techniques may lead to inconsistent physiological information (Tzeng *et al*., 2012; Tzeng & Ainslie, 2013). Additionnally, maneuvers such as the sit-to-stand may provide more realistic data closer to daily life in comparison to data gathered with the thigh-cuff deflation technique. To our knowledge, there is no published report, which examined the dynamic cerebral autoregulatory capacity in endurance athletes using the sit-to-stand maneuver.

It is now recognized that spontaneous TFA has poor signal-to-noise ratio and poor reproducibility compared to TFA performed on forced oscillations (e.g. induced by repeated squat-stand maneuvers) (Smirl *et al*., 2015). Of note, the thigh-cuff deflation technique for quantifying the cerebral autoregulatory index has even worse reproducibility than spontaneous TFA metrics [reviewed in (Smirl *et al*., 2015)]. The above-mentioned issues could explain, at least partly, why findings related to the impact of CRF on dynamic cerebral autoregulatory capacity currently in the literature are equivocal (Lind-Holst *et al*., 2011; Aengevaeren *et al*., 2013; Ichikawa *et al*., 2013). Accordingly, utilizing a multiple assessment approach to study dynamic cerebral autoregulatory capacity within the same individuals is required to improve physiological interpretation. Importantly, this approach should include forced MAP oscillations induced by squat-stand maneuvers to enhance the interpretation of the linear association between arterial blood pressure and cerebral blood velocity (Smirl *et al*., 2015). Importantly, it is possible that the small magnitude and inconsistency of spontaneous MAP oscillations associated with the TFA approach fail to detect subtle abnormalities in dynamic cerebral autoregulatory capacity (Bailey *et al*., 2013a). These modifications may only become obvious when MAP oscillations are forced during sit-to-stand or repeated squat-stand maneuvers.

With the understanding that each dynamic cerebral autoregulatory capacity assessment technique most likely reflects somewhat different components of the cerebral pressure-flow regulation (Tzeng *et al*., 2012), the aims of this study were to address whether CRF affects different metrics using a multiple assessment strategy and hemodynamic stressors (sit-to-stand, TFA on spontaneous and forced MAP and MCAv oscillations). We addressed these aims via a comparative approach between endurance athletes and recreationally active controls and with varying CRF using a correlational approach. In order to examine if the change in dynamic cerebral autoregulatory capacity with CRF represents an adaptive or maladaptive consequence of the trained state, we assessed whether dynamic cerebral autoregulatory capacity was related to orthostatic tolerance in our participants. We hypothesized higher CRF would be associated with a delayed onset of regulation following sit-to-stand but comparable TFA metrics between spontaneous/driven MAP and MCAv oscillations. Finally, the delayed onset of regulation observed with higher CRF would not be associated with reduced orthostatic tolerance.

## MATERIALS AND METHODS

### Participants

Nineteen endurance-trained male athletes (O_2max_ : 55.9 ± 4.9 mL·kg·min^-1^, training history of 5 to 12 hrs·week^-1^ for at least 2 years) and eight inactive/moderately active control participants (O_2max_: 39.2 ± 5.0 mL·kg·min^-1^; p<0.001 vs. athletes, training history <3 hrs·week^-1^) (Table 1) volunteered to participate in this study. All participants provided written informed consent, and the study was approved by the Comité d’éthique de la recherche de l’IUCPQ-Université Laval (CER:20869) according to the principles established by the Declaration of Helsinki. Endurance athletes were trained and competed in a variety of endurance sports, including: cycling (n = 9), triathlon (n = 7), mountain biking (n = 2) and cross-country skiing (n = 1). All participants were free from any medical conditions, demonstrated a normal ECG, and were not taking any medication. This study was part of a larger study characterizing cardiovascular and cerebrovascular function in endurance athletes, and examining high-intensity interval training effects on performance and physiological function.

**Table 1:** Baseline characteristics and resting values between athletes and controls

|  | **Athletes** | **Controls** | **P values** |
| --- | --- | --- | --- |
| **Baseline characteristics** |  |  |  |
| n | 19 | 8 |  |
| Age (y) | 26 ± 5 | 31 ± 4 | 0.07 |
| Height (m) | 1.77 ± 0.08 | 1.80 ± 0.10 | 0.47 |
| Body mass (kg) | 72.6 ± 9.5 | 84.1 ± 12.2 | 0.02 |
| Body mass index (kg/m<sup>2</sup>) | **23 ± 2** | **26 ± 3** | **0.01** |
| Maximal oxygen uptake (ml/kg min<sup>-1</sup>) | **55.9 ± 4.9** | **39.2 ± 5.0** | **<0.001** |
| **Resting values** |  |  |  |
| n | 19 | 8 |  |
| Heart rate (bpm) | 54 ± 8 | 58 ± 5 | 0.20 |
| Mean arterial pressure (mmHg) | 73 ± 10 | 74 ± 10 | 0.92 |
| Middle cerebral artery mean flow velocity (cm/s) | 63.9 ± 11.7 | 65.5 ± 14.8 | 0.75 |
| Cerebrovascular conductance index (cm/s/mmHg) | 0.88 ± 0.14 | 0.91 ± 0.23 | 0.72 |

Values are mean ± standard deviation

### Experimental design

Participants reported to the laboratory on two separate occasions to perform: 1) an incremental cycling test to determine O_2max_, and 2) anthropometric and resting measurements as well as an evaluation of cerebrovascular function and orthostatic tolerance (Figure 1). Participants were asked to refrain from exercise training for at least 12 h and to avoid alcohol and caffeine consumption for 24 h before each visit. All sessions were performed in the same order for all participants and there was at least 24 h between testing sessions.

**Figure 1:**
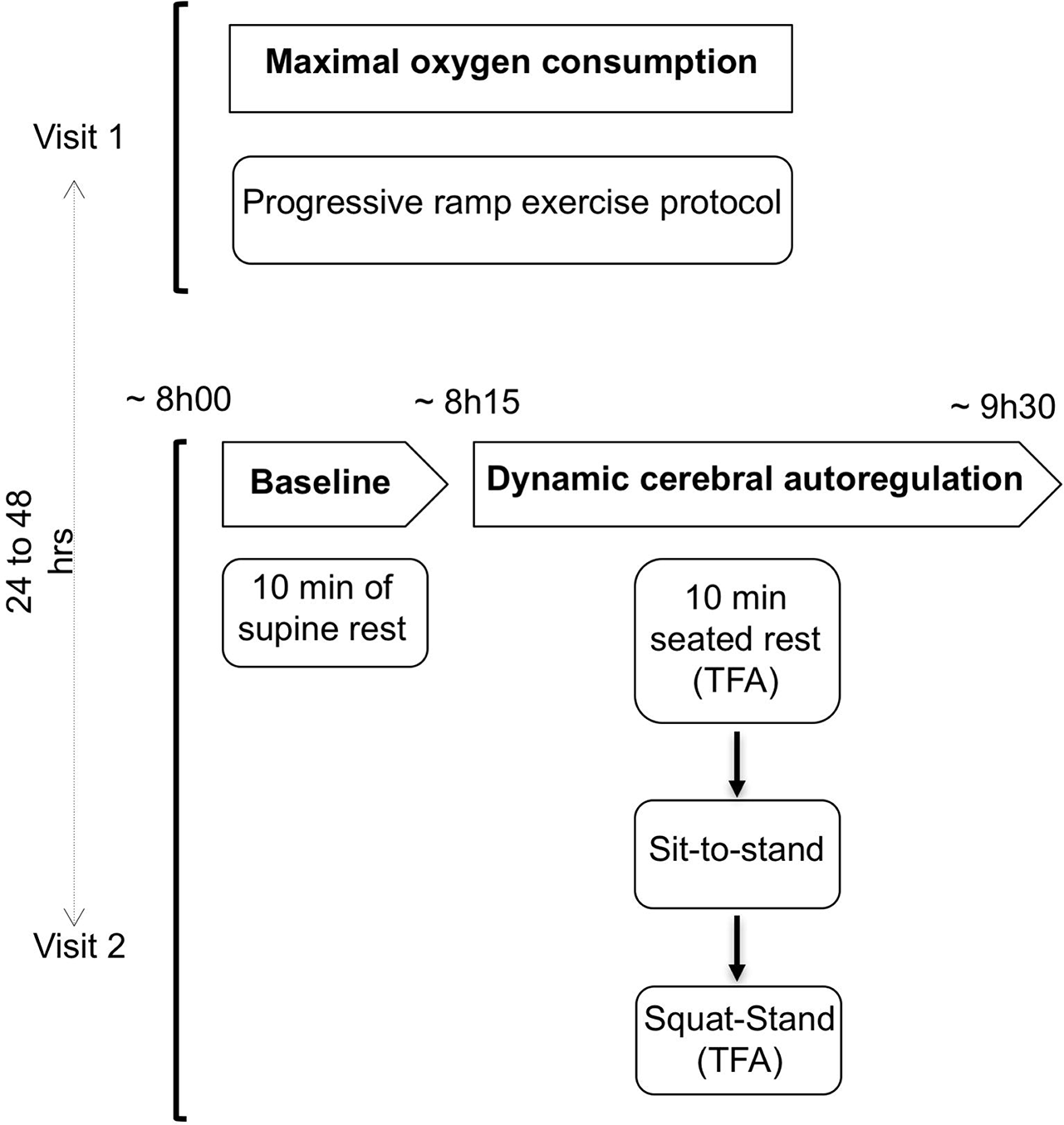
Experimental protocol

### Measurements

#### Heart rate and blood pressure

Heart rate was measured using a 3-lead electrocardiograph (ECG). Beat-to-beat blood pressure (BP) was measured by the volume-clamp method using a finger cuff (Nexfin, Edwards Lifesciences, Ontario, Canada). The cuff was placed on the right middle finger and referenced to the level of the heart using a height correct unit for blood pressure correction. MAP was obtained by integration of the pressure curve divided by the duration of the cardiac cycle. This method has been shown to reliably measure the dynamic changes in beat-to-beat BP which correlate well with the intra-arterial BP recordings and can be used to describe the dynamic relationship between BP and cerebral blood velocity (Omboni *et al*., 1993; Sammons *et al*., 2007).

#### Cerebral blood flow

Cerebral blood flow was estimated by monitoring MCAv via a 2-MHz pulsed transcranial Doppler ultrasound (Doppler Box; Compumedics DWL USA, Inc. San Juan Capistrano, CA). Identification and location of the left MCA was determined using standardized procedures (Willie *et al*., 2011). The probe was attached to a headset and secured with a custom-made headband and adhesive conductive ultrasonic gel (Tensive, Parker Laboratory, Fairfield, NY, USA) to ensure a stable position and angle of the probe throughout testing.

#### End-tidal partial pressure of carbon dioxide (PetCO_2_)

End-tidal partial pressure of carbon dioxide (PetCO_2_) was measured during squat-stand maneuvers through a breath-by-breath gas analyzer (Breezesuite, MedGraphics Corp., MN, USA) calibrated following manufacturer instructions before each evaluation.

#### Data acquisition

For each evaluation, signals (except PetCO_2_) were analog-to-digital-converted at 1000 Hz via an analog-to-digital converter (Powerlab 16/30 ML880; ADInstruments, Colorado Springs, CO, USA) and stored for subsequent analysis using commercially available software (LabChart version 7.1; ADInstruments). PetCO_2_ was acquired separately with Breeze Suite (MedGraphics Corp., MN, USA), and time-aligned with the subsequent data.

### Maximal oxygen consumption (O_2max_)

O_2max_ was determined during a progressive ramp exercise protocol performed on an electromagnetically braked upright cycle ergometer (Corival, Lode, the Netherlands). The test started with 1 min of unloaded pedalling followed by an incremental ramp protocol (25 or 30 W/min) to volitional exhaustion. Expired air was continuously recorded using a breath-by-breath gas analyzer (Breezesuite, MedGraphics Corp., MN, USA) for determinaton of O_2_, carbon dioxide producton (VCO_2_), and respiratory exchange ratio (RER: VCO_2_/O_2_). Maximal O_2_ was defined as the highest 30-s averaged O_2_, concurrent with a RER ≥ 1.15. For more detail and interpretation of these results on gas exchange analysis during this maximal cycling protocol, refer to (Paquette *et al*., 2016).

### Anthropometric measurements and resting hemodynamics

Stature and body mass were measured in each participant. Resting hemodynamic measurements included MAP (volume-clamp method using a finger cuff), which has been validated against intra-arterial pressure (Martina *et al*., 2012), heart rate (HR; ECG) and mean MCAv (MCAv_mean_) (transcranial Doppler ultrasound), which were continuously monitored on a beat-by-beat basis during 10 min of supine rest. Cerebrovascular conductance index (CVCi; MCAv_mean_ /MAP) was then calculated. The average values of the last 5 min of recording represented the baseline.

### Integrative assessment of cerebrovascular function and orthostatic tolerance

To assess dynamic cerebral autoregulatory capacity to transient changes in MAP across the spectrum of physiological extremes (spontaneous to forced MAP oscillations), and given that there is no gold standard methodology for dynamic cerebral autoregulatory capacity quantification (Tzeng *et al*., 2012), we employed a multiple metric approach as outlined below in the chronological order that they were completed (Figure 1).

#### Spontaneous oscillations in MAP and MCAv

Participants rested in a seated position for 10 min. All systemic and cerebral hemodynamic variables were continuously monitored, and the last 5 min of the data from the quiet rest period was used to characterize the dynamic relationship between MAP and MCAv via TFA under spontaneous conditions (see the “Data analysis and statistical approach” section).

#### Sit-to-stand

Subsequent to the 10 min of seated quiet rest, participants quickly (0-3 s) stood up and maintained this standing position for 5 min without moving. MAP and MCAv_mean_ were continuously monitored during this evaluation of the sit-to-stand response. In addition to the characterization of the dynamic cerebral autoregulatory capacity (see the “Data analysis and statistical approach” section), this technique permitted us to examine the orthostatic tolerance of the participants. A participant was considered intolerant when criteria for initial orthostatic hypotension were reached. Initial orthostatic hypotension was defined as a decrease in systolic blood pressure ≥ 40 mmHg and/or a decrease in diastolic blood pressure ≥ 20 mmHg during the first 15 s of standing (Wieling *et al*., 2007). At the end of the standing position, participants were asked to indicate if they experienced any orthostatic symptoms (dizziness, blurry vision, weakness, etc.) after standing. Furthermore, the sit-to-stand technique was used to focus on two principal phases of the orthostatic response: 1-) the initial response (0–30 s), and 2-) the steady-state response (120–180 s) (Wieling & van Lieshout, 1997). The first phase permitted us to analyze the immediate response following sit-to-stand, while the second phase was used to evaluate the recovery of hemodynamic variables.

#### Repeated squat-stand maneuvers

Following 10 min of quiet rest, squat-stand maneuvers were performed after confirmation of the recovery of baseline hemodynamics following sit-to-stand. Participants started in a standing position then squatted down until the back of their legs attained a ∼90 degree angle. This squat position was maintained for a specific period of time (5 or 10 s – see below), after which they moved to the standing position. Following instructions and practice, participants performed 5-min periods of repeated squat-stand maneuvers at a frequency of 0.05 Hz (10-s squat, 10-s standing) and 0.10 Hz (5-s squat, 5-s standing) as previously described (Smirl *et al*., 2015) with 5 min of standing rest between 0.05 and 0.10 Hz maneuvers. These large oscillations in MAP (∼25-30 mmHg) are extensively buffered by the cerebrovasculature when executed at frequencies within the high-pass filter buffering range (<0.20 Hz). The repeated squat-stand maneuvers augment the signal-to-noise ratio enhancing the reproducibility and interpretability of findings through a physiologically-relevant MAP stimulus to the cerebrovasculature (Smirl *et al*., 2015). The sequence of the squat-stand maneuvers was randomized between participants and each frequency was separated by 5 min of recovery. During these maneuvers, participants were instructed to maintain normal breathing and to avoid Valsalva. The linear aspect of the dynamic MAP-MCAv relationship was characterized via TFA (see the “Data analysis and statistical approach” section). MAP, HR, MCAv_mean_ and PetCO_2_ were continuously monitored during this evaluation. PetCO_2_ data was averaged over each 5-min squat-stand maneuvers.

### Data analysis and statistical approach

#### Cerebrovascular responses to acute hypotension

In order to describe the cerebrovascular responses to acute hypotension following the sit-to-stand technique, we used the following metrics: 1) the time delay before onset of regulation; 2) the change in MCAv_mean_ and MAP from baseline to nadir (absolute: Δ MCAv_mean_, ΔMAP; and relative to baseline: Δ MCAv_mean_ (%), ΔMAP(%)) and 3) the rate of regulation (RoR).

Time delay before onset of regulation is the time lapse between sit-to-stand and the increase in CVCi (Lind-Holst *et al*., 2011). The onset of regulation was considered to have occurred the moment CVCi began increasing constantly without any subsequent transient decrease. This metric was evaluated by two different observers (LL and PB). The fall in MCAv_mean_ and MAP is the difference between baseline MCAv_mean_ or MAP and minimum MCAv_mean_ or MAP recorded after the sit-to-stand. The physiological response to acute hypotension can be divided into two phases (Ogoh *et al*., 2008); Phase I is the time point after sit-to-stand where MCAv_mean_ changes are independent of any arterial baroreflex correction (1 to 7 s after sit-to-stand (van Beek *et al*., 2008; Deegan *et al*., 2009; Sorond *et al*., 2009; Tzeng *et al*., 2014)). Phase II is the time point starting at the onset of arterial baroreflex and continuing for 4 sec (Ogoh *et al*., 2008). Thus, during Phase I, the rate of change in CVCi is directly related to dynamic cerebral autoregulatory capacity, without arterial baroreflex regulation. Accordingly, RoR was calculated during Phase I as an index of dynamic cerebral autoregulatory capacity:

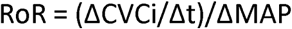

Where ΔCVCi/Δt is the linear regression slope between CVCi and time (t) during Phase I (a 2.5-s interval (Δt) starting after individually determined onset of regulation was taken for the analysis of RoR), and ΔMAP is calculated by subtracting baseline MAP from averaged MAP during Phase I (Aaslid *et al*., 1989; Ogoh *et al*., 2008).

#### Assessment of the dynamic relationship between MAP and MCAv

Beat-to-beat MAP and MCAv signals were spline interpolated and re-sampled at 4 Hz for spectral analysis and TFA based on the Welch algorithm. Each 5-min recording was first subdivided into 5 successive windows that overlapped by 50%. Data within each window were linearly detrended and passed through a Hanning window prior to discrete Fourier transform analysis. For TFA, the cross-spectrum between MAP and MCAv was determined and divided by the MAP auto-spectrum to derive the transfer function coherence, absolute gain (cm/s/mmHg), normalized gain (%/%) and phase (radians) in accordance with the recommendations of the Cerebral Autoregulation Research Network (CARNet)(Claassen *et al*., 2016).

The TFA coherence, gain and phase of the spontaneous oscillations were band averaged across the very low-(0.02-0.07 Hz), low-(0.07-0.2 Hz) and high-frequency (0.2-0.4 Hz) ranges, driven MAP oscillations were sampled at the point estimate of the driven frequency (0.05 and 0.10 Hz). These point estimates were selected as they are in the very low (0.02-0.07 Hz) and low (0.07-0.20 Hz) frequency ranges where cerebral autoregulation is thought to be most operant (Zhang *et al*., 1998). Only the TFA phase and gain values where coherence exceeded 0.50 were included in analysis to ensure the measures were robust for subsequent analysis (Smirl *et al*., 2014). Phase wrap-around was not present when coherence exceed 0.50 in the spontaneous data nor at any of the point-estimate values for squat-stand maneuvers.

#### Statistical analysis

After confirming normal distribution of data using Shapiro-Wilk normality tests, between-group differences were analyzed using independent samples t-tests, with Welch’s correction in the presence of unequal variances between groups. Difference in the prevalence of initial orthostatic hypotension between groups was analyzed using the Fisher’s exact test. Relationships between variables were determined using Pearson product moment or Spearman’s Rho correlations. Statistical significance was established at P < 0.05 and data are expressed as mean ± standard deviation.

## RESULTS

### Participant characteristics and baseline systemic and cerebrovascular hemodynamics

Body mass (p=0.02) and body mass index (p=0.01) were lower in athletes as anticipated. Baseline systemic and cerebrovascular hemodynamics monitored during supine rest were similar between athletes and controls (Table 1). As expected, O_2max_ was higher in athletes (+17 mL·kg·min^-1^; p<0.001) compared to the control group.

### Effects of cardiorespiratory fitness on dynamic cerebral autoregulatory capacity

#### TFA of spontaneous oscillations in MAP and MCAv

MAP and MCAv power spectrum densities during spontaneous oscillations were not different between athletes and controls (Table 2). Athletes and controls also had comparable TFA coherence, gain and normalized gain and phase (all p>0.11).

**Table 2:**
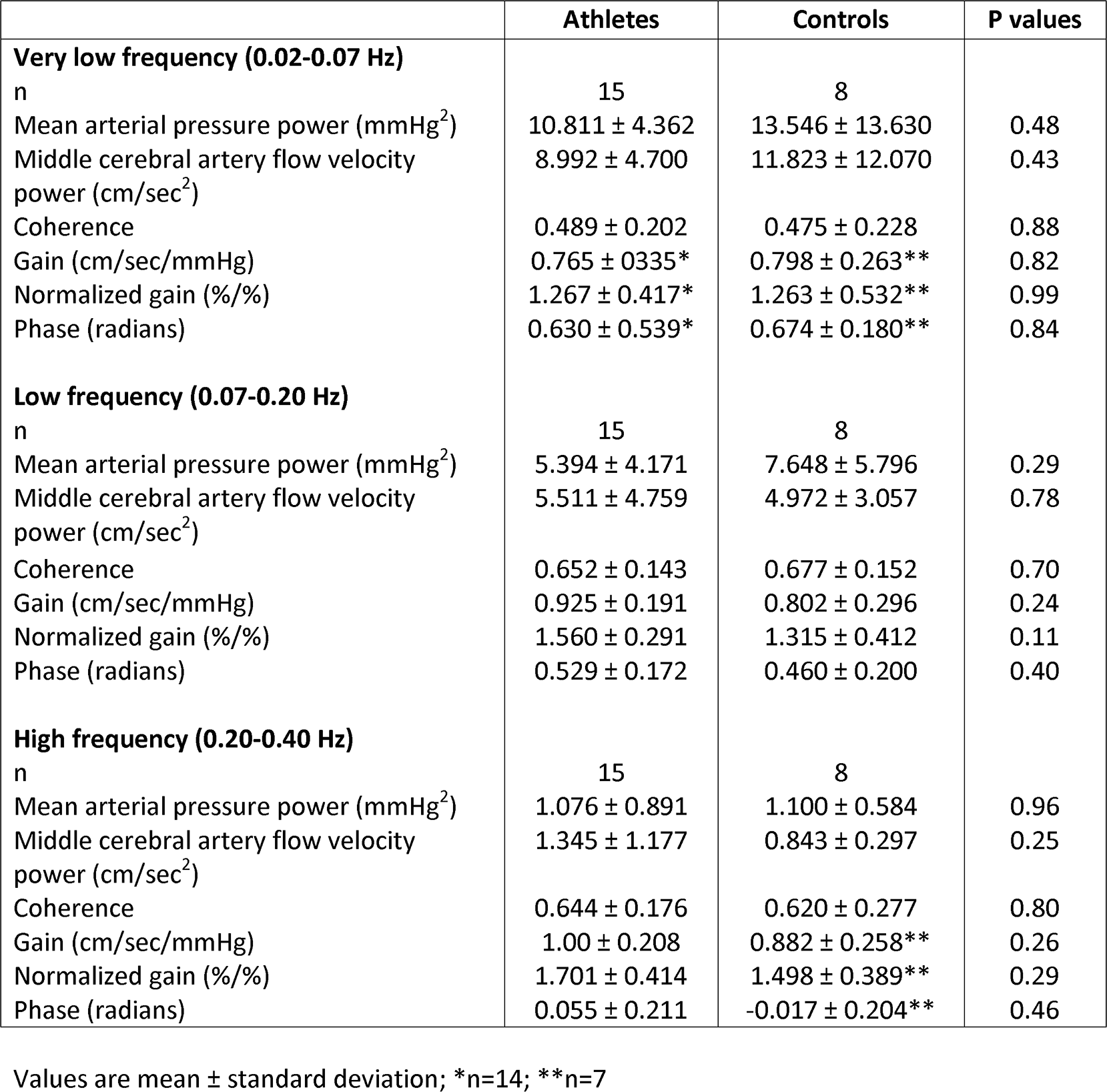
Transfer function analysis of spontaneous oscillations in mean arterial pressure and middle cerebral artery flow velocity

|  | **Athletes** | **Controls** | **P values** |
| --- | --- | --- | --- |
| **Very low frequency (0.02-0.07 Hz)** |  |  |  |
| n | 15 | 8 |  |
| Mean arterial pressure power (mmHg<sup>2</sup>) | 10.811 ± 4.362 | 13.546 ± 13.630 | 0.48 |
| Middle cerebral artery flow velocity power (cm/sec<sup>2</sup>) | 8.992 ± 4.700 | 11.823 ± 12.070 | 0.43 |
| Coherence | 0.489 ± 0.202 | 0.475 ± 0.228 | 0.88 |
| Gain (cm/sec/mmHg) | 0.765 ± 0.335* | 0.798 ± 0.263** | 0.82 |
| Normalized gain (%/%) | 1.267 ± 0.417* | 1.263 ± 0.532** | 0.99 |
| Phase (radians) | 0.630 ± 0.539* | 0.674 ± 0.180** | 0.84 |
| **Low frequency (0.07-0.20 Hz)** |  |  |  |
| n | 15 | 8 |  |
| Mean arterial pressure power (mmHg<sup>2</sup>) | 5.394 ± 4.171 | 7.648 ± 5.796 | 0.29 |
| Middle cerebral artery flow velocity power (cm/sec<sup>2</sup>) | 5.511 ± 4.759 | 4.972 ± 3.057 | 0.78 |
| Coherence | 0.652 ± 0.143 | 0.677 ± 0.152 | 0.70 |
| Gain (cm/sec/mmHg) | 0.925 ± 0.191 | 0.802 ± 0.296 | 0.24 |
| Normalized gain (%/%) | 1.560 ± 0.291 | 1.315 ± 0.412 | 0.11 |
| Phase (radians) | 0.529 ± 0.172 | 0.460 ± 0.200 | 0.40 |
| **High frequency (0.20-0.40 Hz)** |  |  |  |
| n | 15 | 8 |  |
| Mean arterial pressure power (mmHg<sup>2</sup>) | 1.076 ± 0.891 | 1.100 ± 0.584 | 0.96 |
| Middle cerebral artery flow velocity power (cm/sec<sup>2</sup>) | 1.345 ± 1.177 | 0.843 ± 0.297 | 0.25 |
| Coherence | 0.644 ± 0.176 | 0.620 ± 0.277 | 0.80 |
| Gain (cm/sec/mmHg) | 1.00 ± 0.208 | 0.882 ± 0.258** | 0.26 |
| Normalized gain (%/%) | 1.701 ± 0.414 | 1.498 ± 0.389** | 0.29 |
| Phase (radians) | 0.055 ± 0.211 | -0.017 ± 0.204** | 0.46 |

Values are mean ± standard deviation; \*n=14; \*\*n=7

#### Responses to acute hypotension induced by sit-to-stand

MAP (95±10 vs. 97±13 mmHg; p=0.69) and MCAvmean (61±9 vs. 60±13 cms^-1^; p=0.83) were not different between athletes and controls at baseline. Upon standing, decreases in MAP and MCAv_mean_ to their nadir were of similar amplitude in athletes compared to controls (Table 3). There were no group differences in the time taken to reach the nadir for MAP (13±8 vs. 14±9 s; p=0.67) and MCAv_mean_ (7±2 vs. 8±1 s; p=0.54). The time delay before onset of regulation was more than 2-fold longer in athletes (p = 0.03; Table 3 and Figure 2). This metric was inversely correlated with TFA phase during 0.05 Hz (r=-0.499; p=0.02) and 0.1 Hz (r=-0.429; p=0.046) squat-stand, and positively correlated with TFA gain (r=0.501; p=0.02) and ngain (0.462; p=0.04) during 0.1 Hz squat-stand. Following onset of regulation, RoR was similar between groups (p=0.82; Table 3).

**Table 3:** Central and cerebrovascular hemodynamics in response to sit-to-stand

|  | **Athletes** | **Controls** | **P values** |
| --- | --- | --- | --- |
| n | 16 | 7 |  |
| Onset of regulation (s) | **3.1 ± 1.7** | **1.5 ± 1.0** | **0.03** |
| Δ Mean arterial pressure to nadir (mmHg) | -26 ± 10 | -21 ± 5 | 0.11 |
| Δ Middle cerebral artery mean flow velocity to nadir (cm/s) | -15 ± 5 | -11 ± 7 | 0.11 |
| Δ Mean arterial pressure to nadir (%baseline) | -27 ± 10 | -22 ± 5 | 0.25 |
| Δ Middle cerebral artery mean flow velocity to nadir (%baseline) | -25 ± 7 | -18 ± 1 | 0.09 |
| Rate of regulation (s<sup>-1</sup>) | 0.24 ± 0.18 | 0.21 ± 0.22 | 0.82 |

Values are mean ± standard deviation

**Figure 2:**
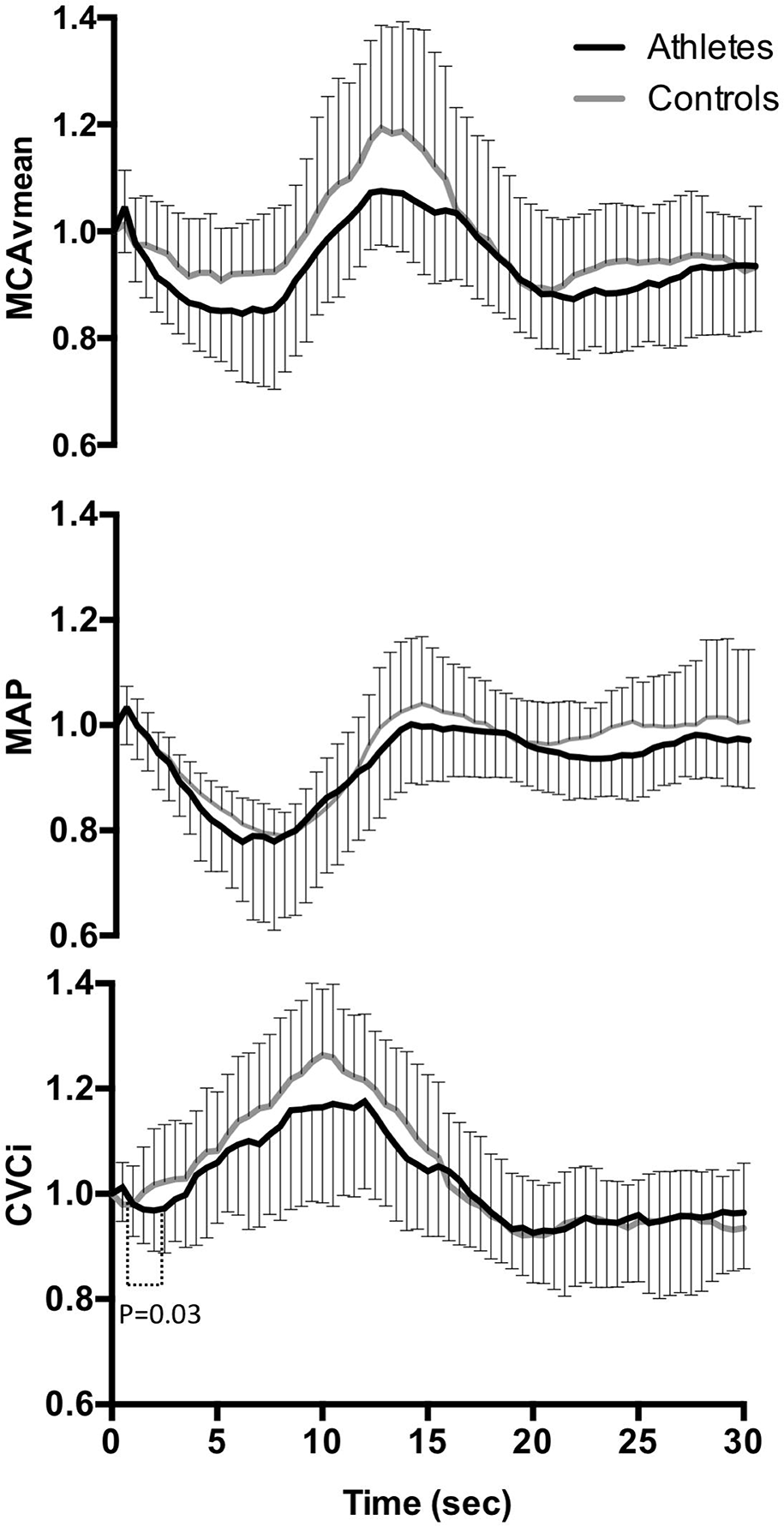
Normalized response of mean arterial pressure (MAP), middle cerebral artery mean flow velocity (MCAv_mean_) and cerebrovascular conductance index (CVCi) in athletes (black line) and controls (grey line) after sit-to-stand. Time 0 indicates the transition from sitting to standing.

The prevalence of initial orthostatic hypotension was comparable between groups (Athletes: 10/16 vs. Controls: 3/7, p=0.65), though none of the participants reported symptoms consistent with neurogenic syncope. None of the participants was classified as orthostatically hypotensive during the steady-state phase (120–180 s) of standing. There were no correlations between metrics of the dynamic cerebral autoregulatory capacity and decreases in MAP or MCAv_mean_ to their nadir upon standing (data not shown).

#### TFA of forced oscillations in MAP and MCAv

MAP and MCAv power spectrum densities during forced oscillations at 0.05 and 0.10 Hz were not different between athletes and controls (Table 4). Absolute TFA gain during 0.10 Hz squat-stand was higher in athletes compared to controls (p = 0.01). Absolute and normalized TFA gain during 0.10 Hz squat-stand were positively correlated with O_2max_ (TFA gain: r = 0.61, p < 0.01; Figure 3A, TFA normalized gain: r = 0.73, p < 0.01; Figure 3B; pooled data), while TFA phase during 0.10 Hz squat-stand was inversely correlated with O_2max_ (r =-0.55; p < 0.001; Figure 3C). All the other metrics were not different between groups (Table 4) and not correlated with O_2max_. PetCO_2_ was comparable between athletes and controls during squat-stand maneuvers performed at 0.05 (43±5 vs. 43±5 mmHg; p=0.95) and 0.10 Hz (43±5 vs. 44±4 mmHg; p=0.75).

**Table 4:** Transfer function analysis of forced oscillations in mean arterial pressure and middle cerebral artery flow velocity using squat-stand maneuvers

|  | **Athletes** | **Controls** | **P values** |
| --- | --- | --- | --- |
| **Squats 0.05 Hz** |  |  |  |
| n | 13 | 7 |  |
| Mean arterial pressure power (mmHg<sup>2</sup>)/Hz | 4558 ± 5632 | 3514 ± 4244 | 0.67 |
| Middle cerebral artery flow velocity power (cm/sec<sup>2</sup>)/Hz | 9351 ± 12072 | 7780 ± 7110 | 0.76 |
| Coherence | 0.975 ± 0.033 | 0.940 ± 0.070 | 0.24 |
| Gain (cm/sec/mmHg) | 0.675 ± 0.185 | 0.553 ± 0.202 | 0.19 |
| Normalized gain (%/%) | 1.126 ± 0.300 | 0.951 ± 0.273 | 0.22 |
| Phase (radians) | 0.786 ± 0.252 | 0.965 ± 0.340 | 0.16 |
| **Squats 0.10 Hz** |  |  |  |
| n | 15 | 7 |  |
| Mean arterial pressure power (mmHg<sup>2</sup>)/Hz | 7523 ± 5104 | 5863 ± 3482 | 0.45 |
| Middle cerebral artery flow velocity power (cm/sec<sup>2</sup>)/Hz | 10712 ± 8847 | 11634 ± 6165 | 0.81 |
| Coherence | 0.995 ± 0.008 | 0.994 ± 0.008 | 0.78 |
| Gain (cm/sec/mmHg) | **0.871 ± 0.162** | **0.686 ± 0.116** | **0.01** |
| Normalized gain (%/%) | 1.431 ± 0.294 | 1.216 ± 0.189 | 0.09 |
| Phase (radians) | 0.541 ± 0.230 | 0.438 ± 0.238 | 0.31 |

**Figure 3:**
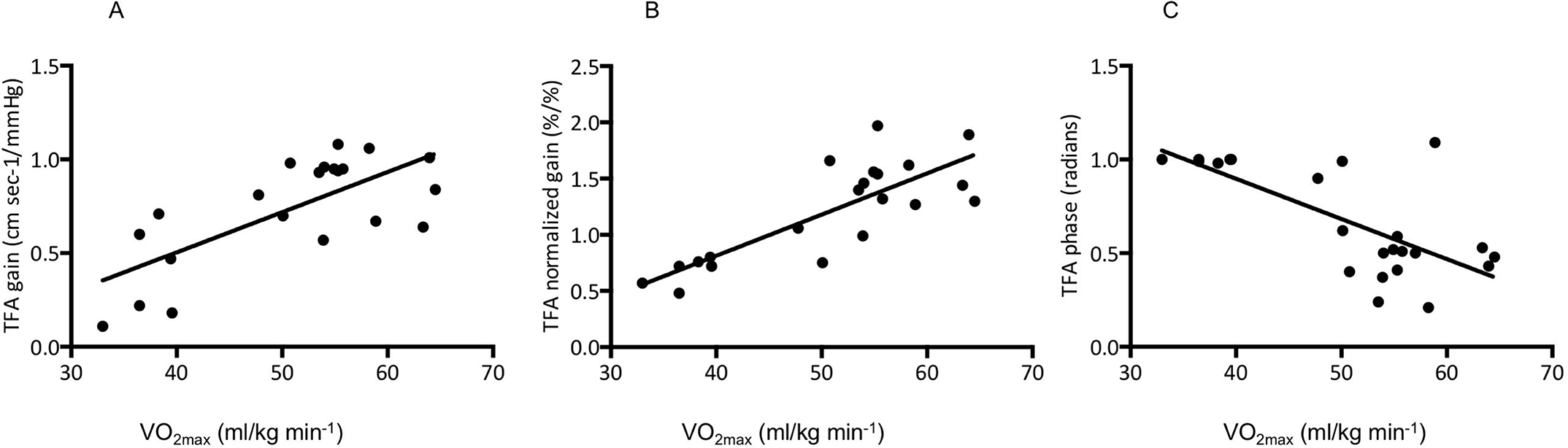
Relationships between maximal oxygen uptake (O_2max_) and TFA gain (Panel A), normalized gain (Panel B) and phase (Panel C) during 0.10 Hz squat-stand. TFA: transfer function analysis

## DISCUSSION

The key findings from this study were that elevated CRF was associated with 1) delayed onset of regulation in response to transient hypotension induced by sit-to-stand, without affecting its response rate (i.e. RoR) and, 2) augmented absolute TFA gain during 0.10 Hz squat-stand. However, TFA metrics during spontaneous MAP oscillations were comparable between groups. Collectively, these results indicate an intact ability of the cerebral vasculature of trained individuals to react to spontaneous MAP oscillations, but a diminished capability to buffer rapid and large MAP changes. Importantly, these findings suggest that we may need to challenge the system in the trained stated (through forced MAP oscillations) to reveal subtle modifications in the dynamic cerebral autoregulatory capacity. In addition, since higher CRF was not associated with a greater prevalence of initial orthostatic hypotension, and that the delayed onset of regulation or augmented TFA gain during 0.10 Hz squat-stand failed to affect orthostatic tolerance, these variations in the dynamic cerebral autoregulatory capacity do not translate into orthostatic intolerance in healthy young male participants. These observations suggest that dynamic cerebral autoregulatory capacity metrics with forced oscillations in MAP is a more sensitive biomarker of impairment compared to the hypotensive challenge used in this study. Considering the differences in the autoregulatory stimulus between each technique, the multimodal approach applied in this study was able to identify subtle differences between these populations, which may have been overlooked if a singular approach was applied (Subudhi *et al*., 2015; Smirl *et al*., 2016).

### Resting cerebral hemodynamics

The literature related to the cerebrovascular benefits of aerobic exercise training is limited and conflicting. Research to date has shown regular exercise is associated with an attenuated decline in cerebral blood velocity with normal aging (Lautenschlager *et al*., 2012), while regular endurance exercise is associated with increased cerebral blood velocity (Ainslie *et al*., 2008) and cerebrovascular reactivity to carbon dioxide (Bailey *et al*., 2013b; Barnes *et al*., 2013). Interestingly, Brugniaux *et al*. (Brugniaux *et al*., 2014) showed the fitness effect for cerebral blood velocity was only apparent during incremental cycling exercise, but not at rest. With this in mind, reduced TFA phase at 0.10 Hz and elevated TFA normalized gain at 0.05 Hz have also been reported with driven oscillations in MAP during moderate exercise with old vs. young adults, while these values were also comparable at rest (Smirl *et al*., 2016). Overall, these findings indicate that cerebrovascular benefits associated with higher CRF may only be discernable when the cerebrovasculature’s regulatory capacity is challenged (Bailey *et al*., 2013a). Our findings of comparable baseline MCAv_mean_ and CVCi between athletes and controls support this notion, albeit within a relatively young and healthy adult population.

### Impact of cardiorespiratory fitness on dynamic cerebral autoregulatory capacity

The influence of CRF on dynamic cerebral autoregulatory capacity remains ambiguous and poorly understood. Lind-Holst *et al*. (Lind-Holst *et al*., 2011) reported a delayed onset of regulation following transient hypotension induced by thigh-cuff deflation and increased TFA normalized gain of spontaneous oscillations in MAP and MCAv in endurance athletes. In contrast, similar dynamic cerebral autoregulatory capacity between young athletes and controls has been reported following thigh-cuff deflation and via TFA of spontaneous oscillations in MAP and MCAv (Ichikawa *et al*., 2013). Further, Aengevaeren *et al*. (Aengevaeren *et al*., 2013) did not observe a difference in dynamic cerebral autoregulatory capacity, characterized by TFA of spontaneous oscillations in MAP and MCAv as well as repeated sit-to-stand, in Masters athletes compared to elderly controls. Such discrepancies may be a consequence, at least partly, of methodological differences in the assessment of dynamic cerebral autoregulatory capacity (Tzeng *et al*., 2012; Smirl *et al*., 2015).

Most metrics of dynamic cerebral autoregulatory capacity appear unrelated to each other or show only weak/moderate correlations (Tzeng *et al*., 2012). As there exists no gold standard to examine dynamic cerebral autoregulatory capacity as a whole entity, several investigators strongly suggest the use of a multiple assessment approach, such as utilized in our study, to improve physiological interpretation of dynamic cerebral autoregulatory capacity (Tzeng *et al*., 2012; Subudhi *et al*., 2015). In support of the findings from Lind-Holst *et al*. (Lind-Holst *et al*., 2011), but with the utilization of a maneuver which represents a physiological stress closer to daily life in comparison to the thigh-cuff deflation technique, we observed that CRF was associated with nearly a 2-s delayed onset of regulation following transient hypotension induced by sit-to-stand (Table 3). However, and similarly to what has been reported by others (Lind-Holst *et al*., 2011), CRF did not influence RoR. Overall, these observations indicate that CRF may affect the latency of cerebrovascular counter-regulation in response to transient hypotension, without disturbing the cerebral autoregulatory control integrity. The physiological and/or clinical significance of a delayed onset of regulation in endurance athletes, or in any other population, remains to be determined. Hypothetically, a postponed initiation of autoregulation in response to an abrupt decrease in ABP could extend the time period associated with pressure-passive cerebral blood velocity without counter-regulation, which in turn would increase to risk of pre-syncope, especially in individuals with an affected cerebral autoregulatory control integrity.

We did not observe any significant difference in TFA metrics during spontaneous oscillations in MAP and MCAv between athletes and controls as reported by some (Aengevaeren *et al*., 2013; Ichikawa *et al*., 2013), but not all (Lind-Holst *et al*., 2011). Interestingly, a higher absolute TFA gain became apparent in athletes during driven oscillations at 0.10 Hz for the squat-stand protocol. These results indicate the cerebral vasculature of individuals with higher CRF has an intact ability to regulate spontaneous oscillations in MAP, but becomes less capable of dampening efficiently large and rapid changes in MAP. These findings could be due to the cardiac baroreceptors aiding in the regulation of MAP for the slower driven oscillations. Indeed, cardiac baroreceptors tend to engage and counter regulate at ∼7 s (Aaslid *et al*., 1989), which is therefore effective for 0.05 Hz, whereas it does not have time to counter-regulate at 0.10 Hz. Also, 0.10 Hz oscillations are associated with Mayer waves, representing an index of sympathetic activity in the peripheral vasculature, which could play a role in the frequency dependent nature of this response (Hyndman *et al*., 1971; Preiss & Polosa, 1974). Contradictory results are present in the literature regarding the influence of CRF on TFA gain. For example, higher (Lind-Holst *et al*., 2011), lower (Ichikawa *et al*., 2013) or similar (Aengevaeren *et al*., 2013) TFA gain has been reported in athletes compared to untrained controls. Differences in body position during evaluation, age of participants, the method used to characterize this metric, as well as fitness levels may all contribute, to some extent, to these discrepancies. Importantly, taking into consideration that spontaneous TFA has rather poor signal-to-noise ratio and reproducibility compared to TFA on driven oscillations (Smirl *et al*., 2015), further research is necessary to better understand whether the changes measured via TFA of spontaneous oscillations in MAP and MCAv reflect a physiologically significant modification in the ability of the cerebral vasculature to regulate MAP. The opposing conclusions in regards to the influence of CRF on dynamic cerebral autoregulatory capacity show the importance of characterizing the latter with multiple metrics, along with the inclusion of various frequencies for repeat driven data methods such as squat-stand manoeuvers, in order to take into account the complete range of physiological information instead of focusing on specific metrics (Tzeng *et al*., 2012; Tzeng & Ainslie, 2013). In addition, the direction of MAP changes should also be considered when studying the dynamic cerebral autoregulatory capacity, as the brain seems to be better adapted to dampen transient hypertension compared to transient hypotension (Tzeng *et al*., 2010; Numan *et al*., 2014; Brassard *et al*., 2017). Future studies to determine how CRF affects this hysteresis phenomenon are needed.

### Variations in the dynamic cerebral autoregulatory capacity in the trained state: Adaptation or maladaptation?

CRF is accepted as an important indicator of cardiovascular health (Ross *et al*., 2016). In addition, elevated CRF is associated with higher resting and exercising cerebral blood velocity across the aging spectrum (Ainslie *et al*., 2008; Brugniaux *et al*., 2014), and cerebrovascular reactivity to carbon dioxide (Bailey *et al*., 2013b). Accordingly, the slower dynamic cerebral autoregulatory capacity in response to transient hypotension as well as subtle changes in the cerebral pressure-flow relationship during squat-stand manoeuvers with elevated CRF reported in this study may seem counter-intuitive, since this would place the brain in a disadvantageous position during hemodynamic stress of large amplitude. On the other hand, our findings may require a re-appraisal of what we expect of the cerebral circulation in response to clear cardiorespiratory adaptation. Of interest, we examined whether these changes in the dynamic cerebral autoregulatory capacity represented adaptive or maladaptive consequences of the trained state by studying relations between dynamic cerebral autoregulatory capacity metrics and orthostatic tolerance in our participants. The prevalence of initial orthostatic hypotension was comparable between athletes and controls, and we did not observe any correlations between dynamic cerebral autoregulatory capacity metrics and reductions in MAP and MCAv_mean_ during orthostatic stress in our participants. Therefore, our findings indicate the functional impairment in dynamic cerebral autoregulatory capacity accompanying higher CRF does not seem to be associated with, nor is critical enough to worsen, orthostatic tolerance. As such, we suggest this functional impairment represents an adaptive, instead of a maladaptive, consequence of the trained state.

This being acknowledged, endurance athletes are exposed, albeit intermittently, to aggressive oscillations in different types of stresses, including shear stress, hypocapnia and hypercapnea, sympathetic nervous activity, inflammation-oxidation-nitrosylation, and blood volume. Accordingly, their cerebral vasculature is intermittently exposed to a very different hemodynamic and metabolic stress milieu compared to non-athletes, and the different impact on cerebrovascular function of each of these stressors remains unclear. The lack of influence of the functional impairment in dynamic cerebral autoregulatory capacity on orthostatic tolerance observed in this study could represent a desensitization of their cerebral vasculature due to chronic exposure to stress (Thomas *et al*., 2013).

Still, we cannot rule out the possibility that this functional impairment in dynamic cerebral autoregulatory capacity could be amplified, and eventually affect orthostatic tolerance, in endurance athletes with higher aerobic capacity. In addition, there is likely a confounding variable at play which is the notion these groups are generally young healthy males and the subtle changes reported in the current study would be exacerbated if these measures were performed in older adults or at risk populations (e.g., hypertension, early stage heart failure or acute mild traumatic brain injury).

### Limitations

There are some limitations to our study that need to be acknowledged. Only young male participants have been studied and our findings cannot be generalized to other populations. Although the PetCO_2_ levels reported in the current study are elevated compared to expected values, they were unchanged throughout the experimental conditions and therefore would have had limited if any impact on the current findings. We acknowledge that the small and unequal sample sizes of our study is associated with interpretive limitations, although a retrospective power analysis revealed that we were adequately powered to observe statistical differences at P < 0.05 for the main variables of interest examined in this study. Another limitation of the present study is its cross-sectional design and a longitudinal study is needed to confirm our findings. Only one sit-to-stand was used for the RoR calculation. The inclusion of more than one trial may be necessary in upcoming studies given the poor reproducibility of this metric. Since age was not perfectly matched between groups (although both were relatively ‘young’ compared to typical older cohorts, e.g., >60 years), our findings may have been influenced by this variable. However, previous work indicates that aging has no influence on dynamic cerebral autoregulatory capacity (Carey *et al*., 2000), although differences may be revealed during exercise (Smirl *et al*., 2016). Cerebral blood velocity in the MCA measured with transcranial Doppler ultrasound will represent flow only if the diameter of the MCA remains constant. Changes in MAP and PetCO_2_ have been associated with changes in the diameter of internal carotid artery and MCA (Coverdale *et al*., 2014; Verbree *et al*., 2014; Lewis *et al*., 2015). We consider that the physiological range of variation in MAP and PetCO_2_ (when available) from this study will be associated with a negligible impact on the diameter of the MCA (Ainslie & Hoiland, 2014). However, further research using a multimodal imaging approach (e.g., combination of transcranial and carotid Doppler ultrasound, where possible), with PetCO_2_ recordings for each procedure, is strongly recommended to support our findings.

### Conclusions

These results indicate an intact ability of the cerebral vasculature to react to spontaneous oscillations but an attenuated capability to counter rapid and large changes in MAP in aerobically fit individuals. Considering the subtle differences in the three techniques used in this study, we strongly suggest other investigators to consider employing a multi-metrics approach for the assessment of dynamic cerebral autoregulatory capacity to improve interpretation and understanding of this measure. By using blood pressure stimuli of diverse natures and magnitudes, the subtle differences in the ability of the cerebral vasculature to respond to MAP changes (whether it is within the context of CRF, or any other physiological situation) are more likely to be revealed.

## AUTHOR CONTRIBUTIONS

LL, KR, SI, MP, OLB, SM, SJEL, DMB, JDS and PB made a substantial contribution to the concept or design, acquisition, analysis, or interpretation of data; LL and PB drafted the article; KR, SI, MP, OLB, SM, SJEL, DMB and JDS revised the article critically for important intellectual content; All authors approved the final version of the manuscript and agree to be accountable for all aspects of the work in ensuring that questions related to the accuracy or integrity of any part of the work are appropriately investigated and resolved. All persons designated as authors qualify for authorship, and all those who qualify for authorship are listed

